# Behavior influences range limits and patterns of coexistence across an elevational gradient in tropical bird diversity

**DOI:** 10.1101/528950

**Authors:** Benjamin G Freeman, Joseph A Tobias, Dolph Schluter

**Affiliations:** Biodiversity Research Centre, University of British Columbia, Vancouver, Canada.; Department of Zoology, University of British Columbia, Vancouver, Canada.; Department of Life Sciences, Imperial College London, London, UK.

**Keywords:** behavior, coexistence, competition, elevational gradient, exploitative competition, interference competition, limiting similarity, range limits

## Abstract

Does competition influence patterns of coexistence between closely related taxa? Here we address this basic question in ecology by analyzing patterns of range overlap between related bird species (“sister pairs”) distributed along a Neotropical elevational gradient. We explicitly contrast the behavioral dimension of interspecific competition (interference competition) with similarity in resource acquisition traits (exploitative competition). We find that behavioral interactions are generally important in setting elevational range limits and preventing coexistence of closely related species. Specifically, close relatives that defend year-round territories tend to live in non-overlapping elevational distributions, while close relatives that do not defend territories tend to broadly overlap in distribution. In contrast, neither similarity in beak morphology nor evolutionary relatedness was associated with patterns of range limitation. Our main result is that interference competition can be an important driver of species ranges at the scale of entire diverse assemblages. Consequently, we suggest that behavioral dimensions of the niche should be more broadly incorporated in macroecological studies.

## Introduction

Understanding the factors that limit species’ distributions is a longstanding goal of ecology (Wallace 1876). One profitable approach to studying range limits is to consider the distributions of closely related species that occur within the same region (Connell 1961, Whittaker 1967, Diamond 1973). For example, many previous studies have focused on how competition for shared limiting resources (exploitative competition) can shape the ranges of related taxa, based on the assumption that species efficient at acquiring resources may be able to exclude less efficient competitors (Gause 1934, Hardin 1960, Tilman 1977). An alternative perspective is that behavioral interactions (interference competition) among related species may limit coexistence and thus determine range limits (Grether et al. 2017). However, few studies have explicitly considered the behavioral dimension of interspecific competition, and whether it contributes to patterns of geographical range limitation.

Elevational gradients provide an excellent system to address the degree to which competition limits species’ ranges. Mountain slopes encompass large environmental variation over a short geographic scale, maximizing the number of closely related species that occur within the same region and minimizing the influence of dispersal constraints on species’ distributions. Interspecific competition is a historically popular hypothesis to explain why species live only within small sections of large elevational gradients (Brown 1971, Diamond 1973, Terborgh and Weske 1975). Although the precise mechanism of interspecific competition is seldom investigated, recent behavioral studies have uncovered cases where range limits along mountain slopes are set in part because species defend territories against related and ecologically similar taxa (e.g., Jankowski et al. 2010; Freeman et al. 2016). For example, two species of singing mice (*Scotinomys* spp.) live in distinct elevational zones in Central America, and behavioral trials and removal experiments show that the behaviorally dominant higher elevation species exhibits territorial aggression that prevents the lower elevation species from expanding upslope (Pasch et al. 2013). These examples provide some support for the key role of behavior in limiting coexistence between close relatives—consistent with MacArthur’s (1972) claim that “behavior reduces a chaotic scramble to an orderly contest.”

These previous findings raise two key questions. First, is the effect of competition restricted to scattered case studies, or does it provide a more general explanation of species’ elevational range limits in diverse assemblages? Second, is the mechanism by which competition sets range limits at macroecological scales linked to exploitative competition, or interference competition? These categories of competition are interrelated because aggressive behavioral interactions likely arise as an adaptive response to underlying competition for resources—i.e., interference competition is based on exploitative competition (Schoener 1983).

To address these two questions, we investigated the distributions of closely related species pairs in a diverse avifauna distributed along a well-studied Andes-to-Amazon elevational gradient with high quality distributional data (Patterson et al. 1998, Walker et al. 2006, Dehling et al. 2014). Specifically, we used trait-based and phylogenetic models to investigate the relative importance of resource acquisition traits versus behavioral traits in determining patterns of coexistence among species pairs. We inferred the intensity of exploitative and interference competition between species pairs as follows (see also Figure 1). For exploitative competition, we measured (1) niche divergence in a resource acquisition trait (beak morphology), because species with similar beaks are predicted to compete for resources more so than species with divergent beaks (Grant and Grant 2006, Pfennig and Pfennig 2012, Pigot et al. 2018), and (2) evolutionary relatedness, because closely related species are thought to generally experience greater competition than distantly related species (Cavender-Bares et al. 2009, Pfennig and Pfennig 2012, Price et al. 2014). For interference competition, we measured strength of territorial behavior, which indicates both overall aggression linked to resource defense and the potential for interspecific territoriality (Ulrich et al. 2018). In sum, our comparative study offers one of the first tests of the relative importance of two interrelated mechanisms by which interspecific competition can set range limits along environmental gradients by preventing closely related species from coexisting.

**Figure 1.**
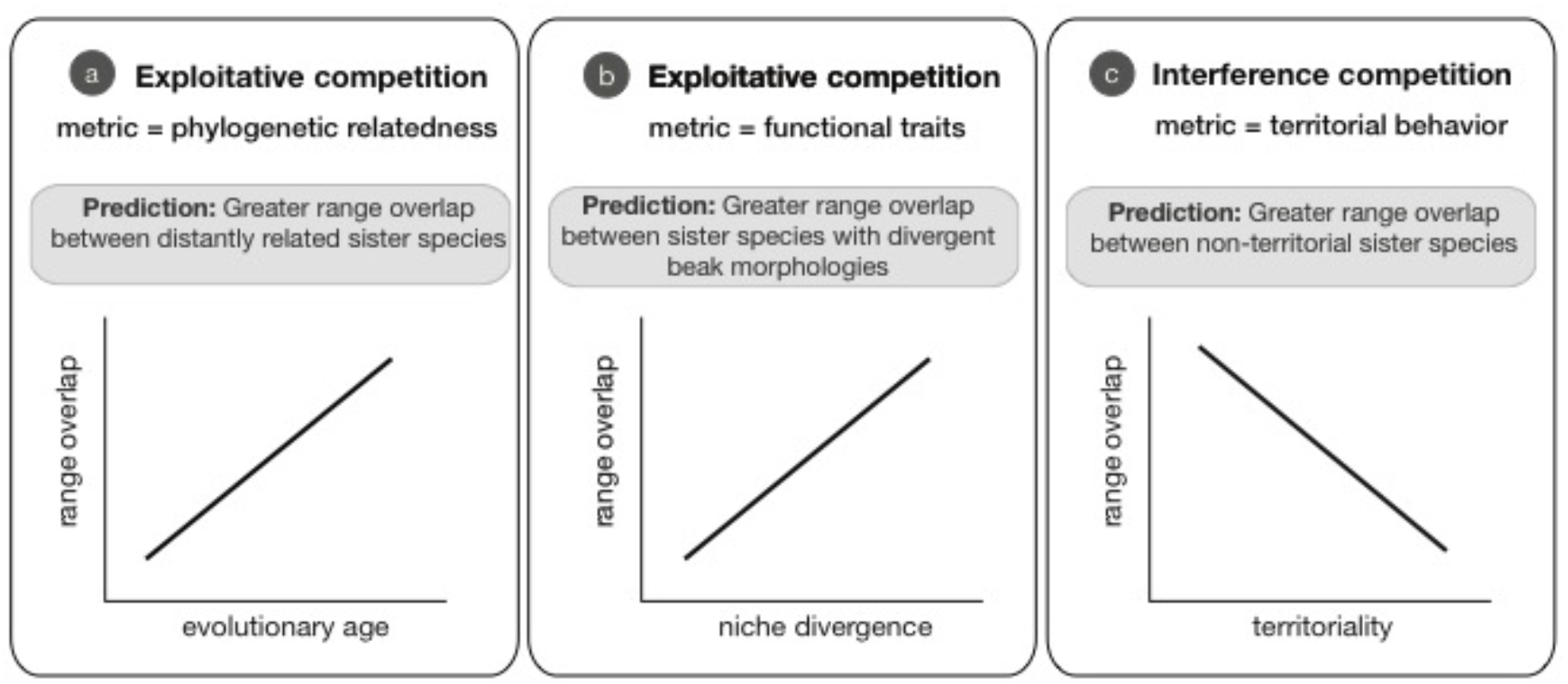
Hypotheses and predictions for patterns of coexistence in sister pairs distributed across an elevational gradient. The hypothesis that exploitative competition influences patterns of coexistence is tested using (a) phylogenetic relatedness and (b) niche divergence in functional traits (beak divergence); the hypothesis that interference competition influences patterns of coexistence is tested using (c) a behavioral trait, territoriality.

## Methods

### Study Region

Our study area is located in the Tropical Andes, home to the greatest concentration of terrestrial biodiversity on Earth (Myers et al. 2000). This “mega” diversity is well illustrated by birds: ~800 bird species occur within our study site—the Manu Transect, a single ~30 km Amazon-to-Andes gradient in southeastern Peru—than across the entirety of North America (Stotz et al. 1996, Walker et al. 2006). Increased levels of biodiversity along mountain slopes arise because high species richness within single elevational zones (alpha-diversity) is coupled with substantial species turnover between elevational zones (beta-diversity). Two examples illustrate the dramatic species turnover along the Manu Transect. First, despite a regional species pool of ~800 resident species, there are only eight species found in both lowland (< 500 m) and high elevation (> 3,000 m) forests (Walker et al. 2006). Second, although this transect spans more than 3,000 m of elevation, the average species inhabits only around one-third of the gradient [elevational breadth = 932 ± 555 m, mean ± standard deviation; *N* = 799 resident species, data from (Walker et al. 2006), omitting non-breeding visitors, species occurring at only a single elevation, species that do not occur along the Manu Transect, and species that live only in agricultural or otherwise highly-modified habitats].

Elevational specialization, in conjunction with high species richness, provides the raw material for our comparative analysis of how competition between closely related species may influence range limits and patterns of coexistence along the Manu Transect. We investigated this question by (1) defining the set of bird species found along the Manu Transect as the regional species pool; (2) defining “sister pairs” within this regional species pool using a molecular phylogeny; (3) measuring evolutionary and ecological variables for each sister pair; and (4) testing the predictions of three hypotheses that attempt to explain why some sister pairs overlap in elevational range along the transect while others do not (Table 1).

**Table 1.**
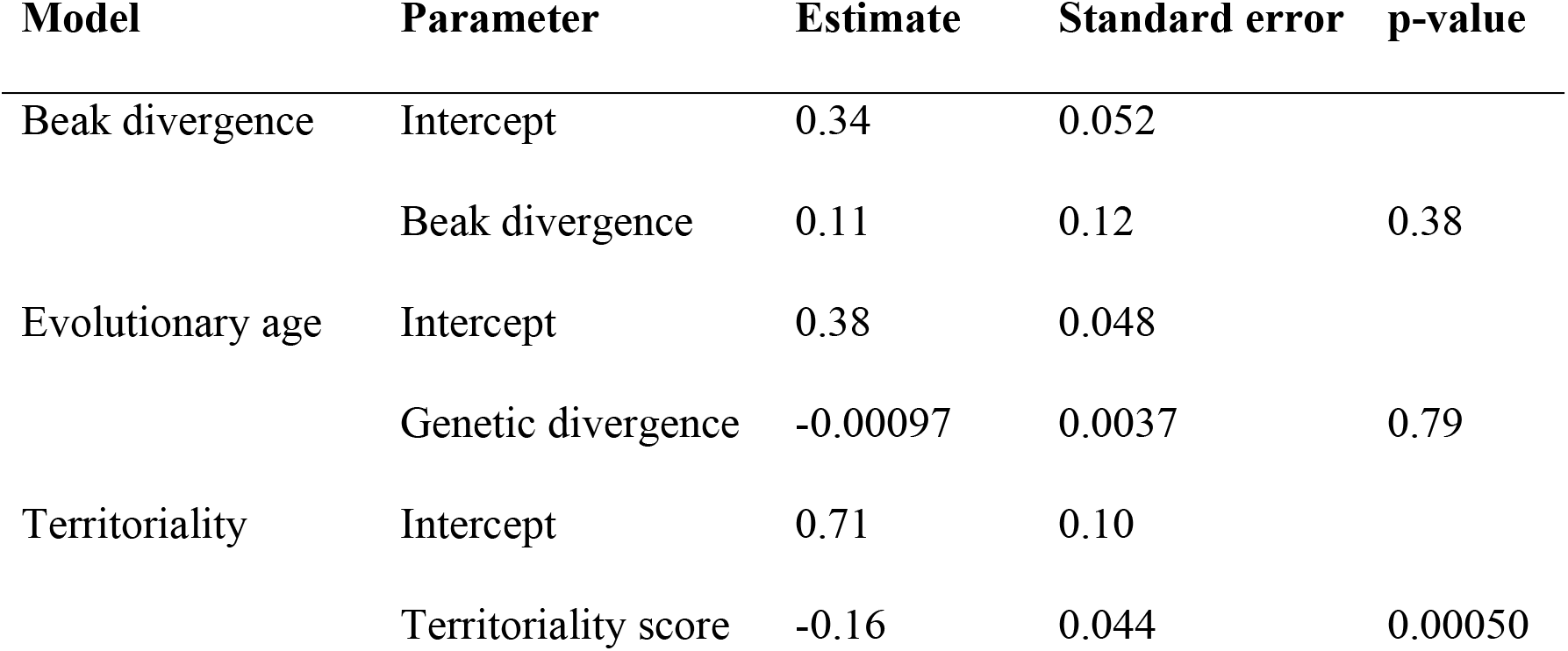
Parameter estimates with standard errors for fixed effects for univariate regression models to predict elevational overlap of sister pairs.

| Model | Parameter | Estimate | Standard error | p-value |
| --- | --- | --- | --- | --- |
| Beak divergence | Intercept | 0.34 | 0.052 |  |
|  | Beak divergence | 0.11 | 0.12 | 0.38 |
| Evolutionary age | Intercept | 0.38 | 0.048 |  |
|  | Genetic divergence | -0.00097 | 0.0037 | 0.79 |
| Territoriality | Intercept | 0.71 | 0.10 |  |
|  | Territoriality score | -0.16 | 0.044 | 0.00050 |

### Defining sister pairs

For our baseline regional species pool, we used a published list of birds recorded in the Manu region (N = 851; Walker et al. 2006). Following Pigot et al. (2016), we removed all non-breeding migrants, all species known from only a single record or elevation, any species not occurring along the best sampled transect (the Manu Transect, sometimes termed the “Manu Road”), and species that live only in agricultural or otherwise highly-modified habitats. This resulted in a final list of 799 species (see Dataset A). We then built a phylogeny of this reduced Manu Transect bird assemblage by subsetting the Manu Transect avifauna from a dated global bird phylogeny (Jetz et al. 2012). Specifically, we downloaded 100 trees from birdtree.org using the Hackett backbone, restricting to taxa included on the basis of genetic information rather than taxonomic inference, and identified the maximum clade credibility (MCC) tree using TreeAnnotator (Rambaut and Drummond 2016). Using this MCC tree, we defined “sister pairs” as lineages that are each other’s closest relatives within the Manu Transect assemblage. Our approach parallels the common usage of “sister pairs” in comparative evolutionary studies, but instead of defining sister taxa among all species on Earth, we restrict ourselves to only those species found in the study transect (a total of 222 sister pairs; see Dataset B). Thus, while some of these “community sisters” are not each other’s closest relatives at global scales, they are each other’s closest relatives within the regional species pool. Before analysis, we restricted our dataset to all sister pairs where at least one species is an upland species with a low elevation limit > 400 m, to avoid the inclusion of sister pairs where both species are restricted to lowland Amazonian forest. A total of 120 sister pairs met this criteria.

### Measuring coexistence

We quantified coexistence as the elevational range overlap between sister pairs. We defined species’ elevational distributions along the Manu Transect using a single published dataset (Walker et al. 2006). This dataset provides elevational limits between 250 m and 4000 m. However, most very high elevation species living along the Manu Transect are coded in this dataset as having an upper elevational limit of 3500 m when in reality they occur up to 4000 m (JAT pers. obs). We therefore extended the upper elevation limit from 3500 m to 4000 m for species that inhabit high elevation puna habitats in this area, using a regional field guide (Schulenberg et al. 2010) and our observations from the field (see Dataset A). We calculated elevational overlap as the percentage of the elevational distribution of the species with the smaller elevational range that overlapped with the larger-ranged species (following Freeman 2015). Thus, sister pairs with non-overlapping elevational ranges had an overlap score of 0, while sister pairs where the range of the smaller-ranged taxa is entirely subsumed within the larger-ranged species had an overlap score of 1. Sister pairs in our dataset varied in the extent of their elevational overlap: 47 sister pairs had zero elevational overlap; 19 had complete elevational overlap; and 54 had intermediate values. Our usage of “coexistence” is consistent with the evolutionary ecology literature that is focused on understanding range limits and spatial variation in species richness (Pigot et al. 2018), but differs from the concept of “stable coexistence” in the theoretical ecology literature, which refers to species’ ability to have a positive long-term growth rate when at low density at a particular site (HilleRisLambers et al. 2012).

### Evolutionary and ecological traits

We defined three evolutionary and ecological traits for each of the 120 sister pairs. First, we calculated beak divergence as the Euclidian distance between species in beak morphospace, following Pigot et al. (2013a). Briefly, we made linear measurements (in mm) of four traits—bill length of the culmen, bill length measured from the nares, bill depth, and bill width—from multiple individuals of each species (mean = 10.55 individuals/species, range = 2 − 107). Measurements were taken from individual birds mist-netted in the field along the Manu Transect, and from specimens stored in museum collections (Trisos et al. 2014, Pigot et al. 2016). We took the log of each of the four traits and ran a principal component analysis (PCA) to generate independent axes of variation in beak morphology. The first two principal components explained nearly the entirety of variation, and were related to overall beak size (PC1, 75.96% of variation) and shape (PC2, 21.20% of variation). We do not present a distinct analysis on body mass divergence because our analysis of beak morphology incorporates both differences in size and in shape. Second, we calculated divergence times from the MCC tree. These divergence times are an estimate of the amount of time (in millions of years) that has elapsed since the two species last shared a common ancestor. Third, we quantified territoriality using a recently published global dataset that classified the territorial defense of all bird species (Tobias et al. 2016). This dataset assigns species to one of three categories: species that do not defend territories (score = 1), species that are weakly or seasonally territorial (score = 2), and species that defend year-round territories (score = 3). See Tobias et al. (2016) for further details on the justification and definition of these categories, and data sources. Because territorial strategies are evolutionarily conserved, most species within a sister pair had identical territoriality scores (29, 36, and 35 sister pairs were designated with scores 1, 2, and 3, respectively). A small number of sister pairs were comprised of constituent taxa with territoriality scores of 1 and 2 (N = 3), or territoriality scores of 2 and 3 (N = 17); in these cases we measured territoriality of the sister pair as the average of the two scores. Importantly, species’ elevational extents were not related to their territoriality score—mean elevational extents for species with territorial scores of 1, 2 and 3 were 1099 m, 1190 m, and 1049 m, respectively.

### Statistical analysis

We conducted all statistical analyses in R (R Development Core Team 2017). We tested our hypotheses (Table 1) by fitting three distinct univariate linear models with elevational overlap as the response variable. We fitted two models to test the importance of exploitative competition in our dataset: a model with beak divergence as the predictor variable, and a model with divergence time as the predictor variable. We also fitted a model with territorial score as the predictor variable to evaluate whether interference competition might explain variation in coexistence in our dataset. We assessed the relative support of the three different univariate models using AIC model selection. We ran (1) ordinary least squares regression models in order to use AIC model selection, and also (2) generalized linear models with family = quasibinomial (link=“logit”) to better approximate the error structure of our data. Last, we also fitted a multiple regression with beak divergence, divergence time and territoriality score as fixed effects.

We tested the influence of phylogenetic non-independence on our data by fitting a phylogenetic generalized least squares (PGLS) multiple regression using the “ape” package (Paradis et al. 2004). The response variable in this model was elevational overlap and fixed effects were territoriality score, divergence time and beak divergence. We estimated evolutionary relationships using a maximum clade credibility tree (implemented in TreeAnnotator (Rambaut and Drummond 2016)) based on 1000 trees downloaded from birdtree.org (Hackett backbone, genetic only) (Jetz et al. 2012), and estimated Pagel’s λ using maximum likelihood. Note that the “traits” in this model (elevational overlap, territoriality score, divergence time and beak divergence) are interactions between two closely related species (the sister pair). In order to run the PGLS, we coded these traits as belonging to one species in the sister pair (i.e., we used a tree with 120 tips, one for each sister pair).

## Results

We report three main results. First, divergence in a resource acquisition trait was minimally related to elevational overlap. We found little support for the prediction that species with different beak morphology were more likely to have overlapping elevational distributions compared to species with similar morphologies (Figure 2b, Table 2). Second, evolutionary age was not associated with coexistence (Figure 2a, Table 2). Third, and in contrast, we found support for the hypothesis that behavior shapes species’ elevational ranges: Sister pairs where both taxa defend year-round territories had much lower elevational overlap than did pairs with weak or absent territoriality (Figure 2c, Table 2, see also Figure 3 for a case example). In the univariate regression model, estimated elevational range overlap was 56% lower for sister pairs that defend year-round territories compared to sister pairs that do not hold territories (estimated range overlaps = 0.25 vs. 0.57, respectively). The univariate model with territoriality score was strongly supported over competing univariate models with beak divergence or evolutionary age (ΔAIC >10; Table S1), and territoriality score was the only significant predictor in a multiple regression model (Table S2). Last, our results are robust to both modeling approaches (Table S3) and phylogenetic non-independence of sister pairs (Table S4).

**Figure 2.**
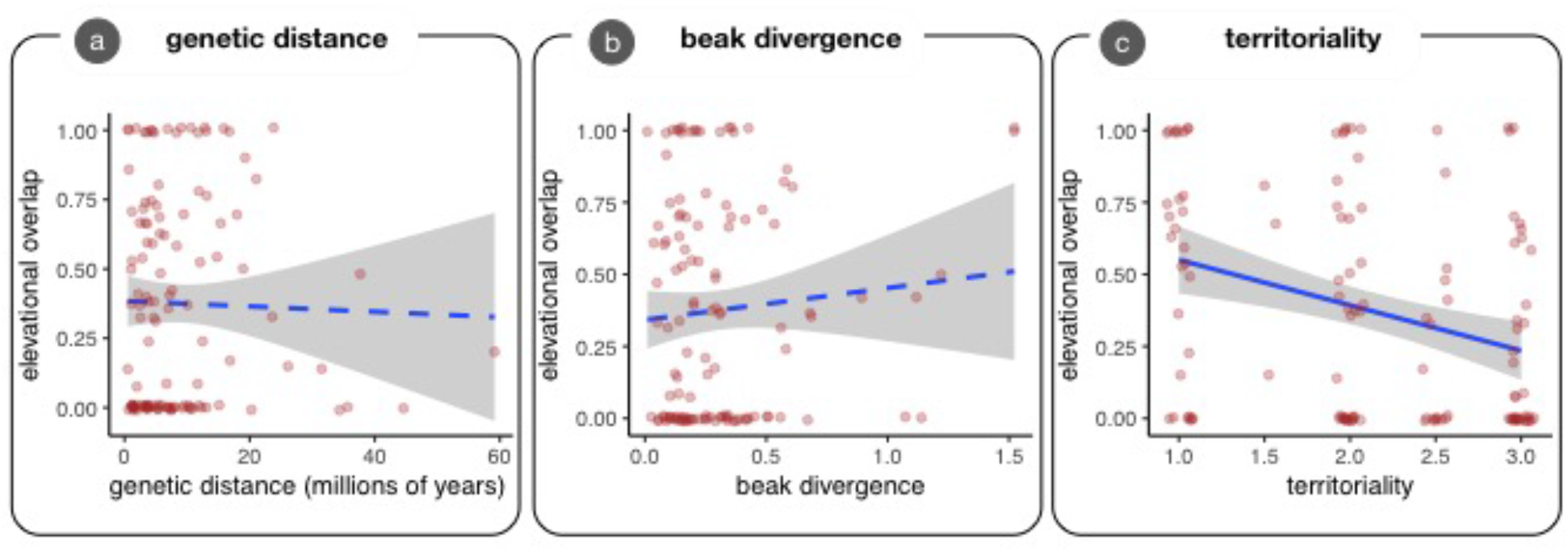
The relationship between elevational overlap of sister pairs and; (a) genetic distance, (b) beak divergence, and (c) territoriality. Best-fit regression lines from univariate regression models are plotted in blue (significant = solid line, non-significant = dashed) with 95% confidence interval shaded in gray. Data are jittered to improve clarity.

**Figure 3.**
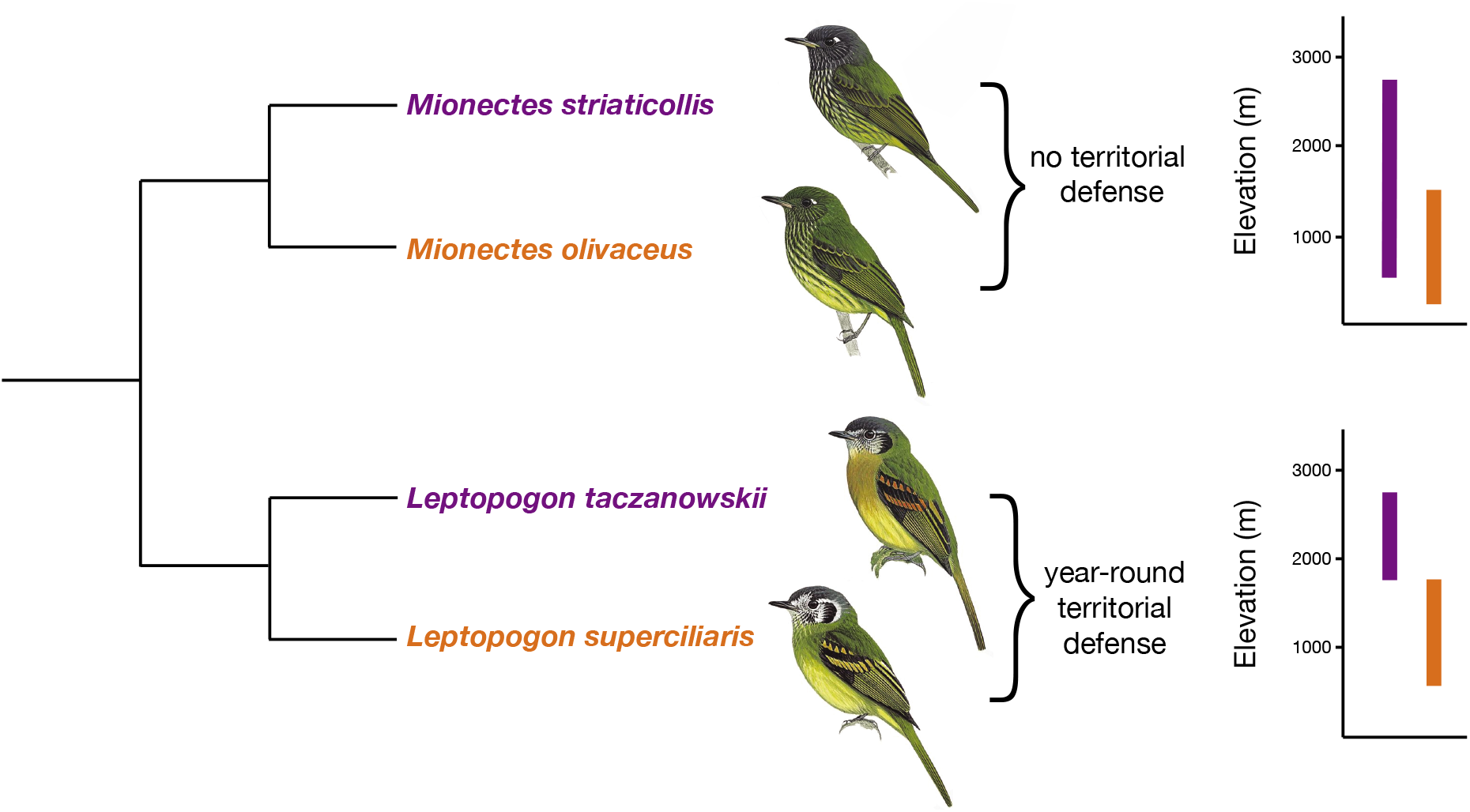
A case example illustrating general patterns. *Mionectes* and *Leptopogon* are sister genera (Miller et al. 2008) that contain morphologically similar sister pairs. *Mionectes* do not defend territories and have broad elevational overlap, while *Leptopogon* defend year-round territories and sharply replace each other at 1800 m. *Mionectes striaticollis* and *olivaceus* are sister taxa (Miller et al. 2008), while *Leptopogon taczanowskii* and *superciliaris* are sister taxa within the Manu avifauna (Winger and Bates 2015). Note that this *Leptopogon* sister pair does not appear in our dataset because genetic sequence data for *Leptopogon* was not available at the time when the molecular phylogeny we used to define sister pairs was built (Jetz et al. 2012). This observation suggests that increased coverage would likely strengthen our findings.

## Discussion

Our main finding is that patterns of coexistence in a diverse Andean avifauna are consistently associated with behavioral traits rather than with morphological trait divergence. That is, patterns of range limitation across an entire bird assemblage appear to be determined more by interference competition than by exploitative competition. Specifically, we find that strength of territoriality is associated with range overlap—closely related species that defend year-round territories tend to live in different elevational zones with minimal overlap, while species that do not defend territories generally overlap in elevational distribution. This general pattern is illustrated by two sister genera of flycatchers, *Mionectes* and *Leptopogon* (Figure 3).

We are at present unable to decisively demonstrate the mechanistic link by which increased territorial behavior leads to reduced range overlap. We hypothesize that the most likely mechanism explaining this pattern is that territorial species often defend their home range against heterospecifics (that tend to be close relatives and ecological competitors), and that interspecific territoriality reduces range overlap between competing species. Supporting this viewpoint, several case examples have demonstrated interspecific territoriality that appears to prevent coexistence of related montane species (Jankowski et al. 2010, Pasch et al. 2013, Freeman et al. 2016).

In contrast, we find no evidence that similarity is related to patterns of coexistence. Hence, limiting similarity (MacArthur and Levins 1967) appears to be absent, at least when considering certain traits (e.g., beak morphology, evolutionary relatedness) at the spatial grain of elevational distributions along a single gradient. We emphasize that the strength of limiting similarity is likely scale-dependent. For example, a previous study also found little signal of limiting similarity when comparing avian assemblages across elevations at our study site using the same traits we studied (Trisos et al. 2014). However, the key result of this previous study was that morphologically and phylogenetically similar taxa seldom overlapped in territories – that is, limiting similarity is strong, but only at the small spatial scale of individual territories (Trisos et al. 2014). At slightly larger scales, limiting similarity also appears to structure bird assemblages in small forest fragments (Ulrich et al. 2018). These findings are consistent with the view that competitive effects are strongest at small spatial scales and decline with increasing spatial scale (e.g., Bullock et al. 2000, Cavender-Bares et al. 2006). However, we note that the dimensions of the niche which we quantified in our study are rather simplistic (e.g., we did not include diet, foraging strata, foraging strategy, or microhabitat), and that patterns consistent with limiting similarity have sometimes been found at biogeographic scales (Pigot and Tobias 2013),

The robustness of our interpretation depends on the validity of our metrics of competition. The metrics we use as proxies for intensity of exploitative competition—morphological and evolutionary similarity—are widely used in the literature, but have been challenged. For example, evolutionary relatedness may not be a reliable proxy for the intensity of competition (Mayfield and Levine 2010), casting doubt on the usefulness of phylogenetic relationships as a proxy for intensity of exploitative competition. In contrast, the assumption that species with similar resource acquisition traits tend to compete more strongly for resources is likely to generally hold. For example, the link between similarity in beak morphology and competition for resources in birds is particularly well supported (e.g., Grant and Grant 2006, Ryan et al. 2007). Perhaps our most important assumption is that the intensity of territorial behavior is a useful metric of interference competition between species. It seems reasonable that intraspecific territorial behavior is a precondition for interspecific territorial behavior—we are not aware of cases where a species defends its territory against heterospecifics but not conspecifics. Nevertheless, further work measuring interspecific territorial defense in the field in tropical taxa would be necessary to test this assumption.

### Implications

To what degree can our results, which apply to a particular assemblage along a particular transect, be generalized to other geographic arenas? We suggest that our primary result—the importance of behavioral interactions to understanding patterns of coexistence—is likely to be of general importance. Interspecific defense of territories has been commonly noted in both tropical and temperate zone birds (Garcia 1983, Robinson and Terborgh 1995, Seddon and Tobias 2010, Losin et al. 2016), as well as in a variety of other vertebrate groups (e.g., Griffis and Jaeger 1998, Pasch et al. 2013). Hence, we hypothesize that territorial interactions may often limit coexistence of close relatives within certain environments (e.g., in this study, different elevational zones). Supporting this conjecture, previously documented cases of interspecific territoriality are often associated with specialization on different microhabitats (Garcia 1983, Robinson and Terborgh 1995, Seddon and Tobias 2010). Further research should investigate whether the common observation that closely related taxa specialize on different habitats in sympatry may be partially driven by behavioral interactions, and whether there are latitudinal trends in such relationships.

### Conclusions

In conclusion, our study highlights that behavioral interactions can be generally important in setting elevational range limits and preventing coexistence of closely related species in a diverse assemblage. This finding adds weight to recent evidence that territorial behavior plays a significant role in structuring tropical montane bird communities (Ulrich et al. 2018). In contrast, we find no direct evidence that limiting similarity shapes distributional limits at the scale of elevational transects, in line with some previous studies (e.g., Trisos et al. 2014). The quest to meaningfully quantify species’ niches has a long history (e.g., Hutchinson 1959). Here we add to this literature by finding that territorial system explains patterns of range limitation better than morphological traits linked to resource acquisition, supporting MacArthur’s (1972) claim that “behavior reduces a chaotic scramble to an orderly contest.” We conclude by reiterating our suggestion that macroecologists interested in explaining large-scale patterns of coexistence ought to consider behavioral dimensions of the niche.

## Acknowledgements

We thank Mark Adams and Hein Van Grouw (Natural History Museum, Tring) and James Van Remsen (Louisiana State University Museum of Natural History) for logistical assistance and access to specimens. We are also grateful to all who contributed morphometric data, including Tom Bregman, Vivien Chua, Bianca Darski, Michael Harvey, Hannah McGregor, Sonia Salazar, Catherine Sheard, and Chris Trisos. This research was supported by postdoctoral fellowships from the Biodiversity Research Centre and Banting Canada (#379958) to BGF, and a Natural Environment Research Council (NE/I028068/1) grant to JAT. Comments from the Schluter lab and Ralf Yorque greatly improved this manuscript. Jonathan Rolland helped write a particularly useful R script, and Ken Thompson helped prepare figures.

## Literature cited

Brown, J. H. 1971. Mechanisms of competitive exclusion between 2 species of chipmunks. Ecology 52:305–311.

Bullock, J. M., R. J. Edwards, P. D. Carey, and R. J. Rose. 2000. Geographical separation of two Ulex species at three spatial scales: does competition limit species’ ranges? Ecography 23:257–271.

Cavender-Bares, J., A. Keen, and B. Miles. 2006. Phylogenetic structure of Floridian plant communities depends on taxonomic and spatial scale. Ecology 87:109–122.

Cavender-Bares, J., K. H. Kozak, P. V. A. Fine, and S. W. Kembel. 2009. The merging of community ecology and phylogenetic biology. Ecology Letters 12:693–715.

Connell, J. H. 1961. The influence of interspecific competition and other factors on the distribution of the barnacle Chthamalus stellatus. Ecology 42:710–723.

Dehling, D. M., S. A. Fritz, T. Töpfer, M. Päckert, P. Estler, K. Böhning-Gaese, and M. Schleuning. 2014. Functional and phylogenetic diversity and assemblage structure of frugivorous birds along an elevational gradient in the tropical Andes. Ecography 37:1047–1055.

Diamond, J. M. 1973. Distributional ecology of New Guinea birds: Recent ecological and biogeographical theories can be tested on the bird communities of New Guinea. Science 179:759–769.

Freeman, B. G. 2015. Competitive Interactions upon Secondary Contact Drive Elevational Divergence in Tropical Birds. The American Naturalist 186:470–479.

Freeman, B. G., A. M. Class Freeman, and W. M. Hochachka. 2016. Asymmetric interspecific aggression in New Guinean songbirds that replace one another along an elevational gradient. Ibis 158:726–737.

Garcia, E. F. J. 1983. An Experimental Test of Competition for Space between Blackcaps Sylvia atricapilla and Garden Warblers Sylvia during the Breeding Season. Journal of Animal Ecology 52:795–805.

Gause, G. F. 1934. The Struggle for Existence. Williams and Wilkins, Baltimore.

Grant, P. R., and B. R. Grant. 2006. Evolution of character displacement in Darwin’s finches. Science 313:224–226.

Grether, G. F., K. S. Peiman, J. A. Tobias, and B. W. Robinson. 2017. Causes and Consequences of Behavioral Interference between Species. Trends in Ecology and Evolution 32:760–772.

Griffis, M., and R. Jaeger. 1998. Competition leads to an extinction-prone species of salamander: Interspecific territoriality in a metapopulation 79:2494–2502.

Hardin, G. 1960. The Competitive Exclusion Principle. Science 131:1292–1297.

HilleRisLambers, J., P. B. Adler, W. S. Harpole, J. M. Levine, and M. M. Mayfield. 2012. Rethinking Community Assembly through the Lens of Coexistence Theory. Annual Review of Ecology, Evolution, and Systematics 43:227–248.

Hutchinson, G. E. 1959. Homage to Santa Rosalia or why are there so many kinds of animals? The American Naturalist 93:145–159.

Jankowski, J. E., S. K. Robinson, and D. J. Levey. 2010. Squeezed at the top: Interspecific aggression may constrain elevational ranges in tropical birds. Ecology 91:1877–1884.

Jetz, W., G. H. Thomas, J. B. Joy, K. Hartmann, and A. O. Mooers. 2012. The global diversity of birds in space and time. Nature 491:444–448.

Losin, N., J. P. Drury, K. S. Peiman, C. Storch, and G. F. Grether. 2016. The ecological and evolutionary stability of interspecific territoriality. Ecology Letters 19:260–267.

MacArthur, R., and R. Levins. 1967. The limiting similarity, convergence, and divergence of coexisting species. American Naturalist 101:377–385.

Mayfield, M. M., and J. M. Levine. 2010. Opposing effects of competitive exclusion on the phylogenetic structure of communities. Ecology Letters 13:1085–1093.

Miller, M. J., E. Bermingham, J. Klicka, P. Escalante, F. S. R. do Amaral, J. T. Weir, and K. Winker. 2008. Out of Amazonia again and again: episodic crossing of the Andes promotes diversification in a lowland forest flycatcher. Proceedings of the Royal Society B: Biological Sciences 275:1133–1142.

Myers, N., R. A. Mittermeier, C. G. Mittermeier, G. A. B. da Fonseca, and J. Kent. 2000. Biodiversity hotspots for conservation priorities. Nature 403:853–858.

Paradis, E., J. Claude, and K. Strimmer. 2004. APE: analyses of phylogenetics and evolution in R language. Bioinformatics 20:289–290.

Pasch, B., B. M. Bolker, and S. M. Phelps. 2013. Interspecific dominance via vocal interactions mediates altitudinal zonation in Neotropical singing mice. The American Naturalist 182:E161–E173.

Patterson, B. D., D. F. Stotz, S. Solari, J. W. Fitzpatrick, and V. Pacheco. 1998. Contrasting patterns of elevational zonation for birds and mammals in the Andes of southeastern Peru. Journal of Biogeography 25:593–607.

Peers, M. J. L., D. H. Thornton, and D. L. Murray. 2013. Evidence for large-scale effects of competition: niche displacement in Canada lynx and bobcat. Proceedings of the Royal Society B: Biological Sciences 280:20132495.

Pfennig, D. W., and K. S. Pfennig. 2012. Evolution’s Wedge: Competition and the Origins of Diversity. University of California Press.

Pigot, A. L., W. Jetz, C. Sheard, and J. A. Tobias. 2018. The macroecological dynamics of species coexistence in birds. Nature Ecology and Evolution 2:1112–1119.

Pigot, A. L., and J. A. Tobias. 2013. Species interactions constrain geographic range expansion over evolutionary time. Ecology Letters 16:330–338.

Pigot, A. L., C. H. Trisos, and J. A. Tobias. 2016. Functional traits reveal the expansion and packing of ecological niche space underlying an elevational diversity gradient in passerine birds. Proceedings of the Royal Society B: Biological Sciences 283:1–9.

Price, T. D., D. M. Hooper, C. D. Buchanan, U. S. Johansson, D. T. Tietze, P. Alström, U. Olsson, M. Ghosh-Harihar, F. Ishtiaq, S. K. Gupta, J. Martens, B. Harr, P. Singh, and D. Mohan. 2014. Niche filling slows the diversification of Himalayan songbirds. Nature 509:222–225.

R Development Core Team. 2017. R: A language and environment for statistical computing. R Foundation for Statistical Computing, Vienna, Austria.

Rambaut, A., and A. Drummond. 2016. TreeAnnotator v.1.8.4.

Robinson, S. K., and J. Terborgh. 1995. Interspecific aggression and habitat selection by Amazonian birds. Journal of Animal Ecology 64:1–11.

Ryan, P. G., P. Bloomer, C. L. Moloney, T. J. Grant, and W. Delport. 2007. Ecological speciation in South Atlantic island finches. Science 315:1420–1423.

Schoener, T. W. 1983. Field experiments on interspecific competition. The American Naturalist 122:240–285.

Schulenberg, T. S., D. F. Stotz, D. F. Lane, J. P. O’Neill, and T. A. Parker. 2010. Birds of Peru. Princeton University Press, Princeton.

Seddon, N., and J. A. Tobias. 2010. Character displacement from the receiver’s perspective: species and mate recognition despite convergent signals in suboscine birds. Proceedings of the Royal Society - Biological sciences 277:2475–2483.

Stotz, D. F., J. W. Fitzpatrick, T. A. Parker, and D. K. Moskovits. 1996. Neotropical birds: ecology and conservation. Cambridge Univ Press, Cambridge, UK.

Terborgh, J., and J. S. Weske. 1975. Role of competition in distribution of Andean birds. Ecology 56:562–576.

Tilman, D. 1977. Resource Competition between Plankton Algae: An Experimental and Theoretical Approach. Ecology 58:338–348.

Tobias, J. A., C. Sheard, N. Seddon, A. Meade, A. J. Cotton, and S. Nakagawa. 2016. Territoriality, Social Bonds, and the Evolution of Communal Signaling in Birds. Frontiers in Ecology and Evolution 4:1–15.

Trisos, C. H., O. L. Petchey, and J. A. Tobias. 2014. Unraveling the interplay of community assembly processes acting on multiple niche axes across spatial scales. The American Naturalist 184:593–608.

Ulrich, W., C. Banks-Leite, G. De Coster, J. C. Habel, H. Matheve, W. D. Newmark, J. A. Tobias, and L. Lens. 2018. Environmentally and behaviourally mediated co-occurrence of functional traits in bird communities of tropical forest fragments. Oikos 127:274–284.

Walker, B., D. F. Stotz, T. Pequeno, and J. W. Fitzpatrick. 2006. Birds of the Manu Biosphere Reserve. Fieldiana: Zoology 110:23–49.

Wallace, A. 1876. The geographical distribution of animals.

Whittaker, R. H. 1967. Gradient analysis of vegetation. Biological Reviews 42:207–264.

Winger, B. M., and J. M. Bates. 2015. The tempo of trait divergence in geographic isolation: avian speciation across the Marañon Valley of Peru. Evolution 69:772–787.

